# CryoEM structure of the *Trypanosoma brucei* flagellum central apparatus

**DOI:** 10.1101/2025.05.29.656921

**Authors:** Saarang Kashyap, Xian Xia, Hui Wang, Michelle Shimogawa, Angela Agnew, Kent Hill, Z. Hong Zhou

## Abstract

Axonemes are cylindrical bundles of microtubule filaments that typically follow a 9+n pattern (where *n* ranges from 0 to 4). However, variations exist across species and cell types, including architectures with fewer (e.g., 3+0, 6+0) or more than nine doublet microtubules (e.g., 9+9+0, 9+9+3), reflecting diverse structural adaptations of cilia and flagella in eukaryotes. *Trypanosoma brucei*, the causative agent of African trypanosomiasis, relies on its single 9+2 flagellum to navigate through environments within the mammalian host and insect vector. Central to the *T. brucei* flagellum’s function is a canonical central apparatus (CA), composed of two—C1 and C2— singlet microtubules, which regulates flagellar beating and ensures efficient movement. Despite its crucial mechanoregulatory role in flagellar beating, the molecular structure and interactions governing *T. brucei* CA assembly and function remain poorly understood. In this study, we employed cryogenic electron microscopy (cryoEM) to uncover structural details of the *T. brucei* CA. We identified conserved and stably C1/C2-associated protein densities, including the armadillo repeat protein PF16, which serves as a structural scaffold critical for CA assembly and axonemal asymmetry. Our analysis also revealed pronounced molecular flexibility of the CA and uncovered *T. brucei*-specific densities, suggesting lineage-specific adaptations for parasite motility. These findings provide critical insights into the structural foundations of *T. brucei* motility. They also highlight potential therapeutic targets to disrupt the parasite’s ability to cause disease, offering new avenues for the treatment of African trypanosomiasis. Comparison of CAs in this canonical 9+2 axoneme and non-canonical 9+n axonemes offers general insights into the assembly and diverse functions of CAs across a wide range of species.

**Significance:** The flagellum of *Trypanosoma brucei,* the parasite causing African trypanosomiasis, drives motility essential for host infection and disease transmission. Our cryogenic electron microscopy study reveals the molecular architecture of its central apparatus, identifying PF16 as a key scaffold that stabilizes axonemal asymmetry and imparts flexibility critical for flagellar beating. We uncover conserved proteins and *T. brucei*- specific adaptations, highlighting evolutionary divergence underlying the parasite’s distinctive motility. These structural results offer insights about how the central apparatus regulates *T. brucei*’s bihelical motion and unveil potential therapeutic targets to disrupt flagellar function and combat African trypanosomiasis. Comparison among the central apparatus of canonical and non-canonical axonemes allow us to suggest broader principles underlying their central roles in ciliary functions across eukaryotes.

## Introduction

The axoneme is an intricate microtubule-based structure that provides the core structural framework of cilia and flagella, organelles critical for cellular motility and signaling (1). Axonemes typically exhibit a canonical “9+2” configuration, consisting of nine outer doublet microtubules (DMTs) surrounding a central microtubule structure. The central microtubule structure consists of a pair of microtubules, known as the central apparatus, extensively decorated by a sheath of proteins. The central apparatus plays a pivotal role in resisting compressive forces of bending DMTs and regulating the coordinated beating of cilia and flagella by interacting with DMT-associated complexes like dynein arms, the molecular motors that drive bending of the flagellum, and radial spokes, which transmit regulatory signals from the central microtubules to dynein motors (2–4) This dynamic architecture maintains axoneme stability while ensuring precise control of ciliary and flagellar function, making the CA essential for processes like sperm propulsion, mucus clearance, embryonic development, and motility of unicellular organisms, including pathogens (5, 6).

The atomic-level understanding of the central microtubules within the axoneme has advanced significantly in recent years, thanks to advancement in cryogenic electron microscopy (cryoEM) and tomography (cryoET). Early studies, such as those on *Chlamydomonas* flagella, mapped central apparatus-associated proteins like hydin and CPC1 at a 16-nm periodicity (2) (7). In 2022, two pivotal studies elucidated the near-atomic resolution CA structures of the axoneme from the green alga, *Chlamydomonas reinhardtii* (8, 9). Han et al. revealed a sophisticated assembly of 45 distinct proteins forming the central apparatus (both microtubule inner and outer proteins) with an overall 32 nm periodicity that coordinates dynein-driven motility. Concurrently, Gui and collaborators reported a complementary structure, detailing the molecular bridges and flexible linkages between the C1 and C2 microtubules and identifying key regulatory proteins that stabilize the apparatus and modulate ciliary beating (9). However, evolutionary divergence has led to substantial variation in CA structure and mechanics across eukaryotes, underscoring the importance of determining CA architecture in a broader range of organisms. Notably, while the CP rotates during beating in *Chlamydomonas* and some other flagellates, it remains static in *Trypanosoma brucei* and several other protists and metazoans, highlighting key differences in how axonemal asymmetry and motility are regulated across species.(10) (11). Together, these studies offer a comprehensive molecular blueprint of the CA, advancing our understanding of its role in flagellar function.

*Trypanosoma brucei*, a parasitic protozoan responsible for African trypanosomiasis, relies on its flagellum for motility and survival (12, 13). The parasite flagellum features a canonical 9+2 axoneme with a central pair of microtubules, critical to its bihelical motion (13) (14). CryoET has resolved the 96-nm axonemal repeat in *T. brucei*, identifying lineage-specific inter-doublet linkages and microtubule inner proteins (MIPs) (15) alongside a lineage-specific extra-axonemal paraflagellar rod (16, 17), that support the unique motility of the parasite. More recently, cryoEM analysis of individual DMTs at ∼3 Å resolution resolved *T. brucei* MIPs and associated dynein and radial spoke proteins with atomic details, elucidating the molecular mechanisms underlying axonemal motility (18, 19). These structural features suggest evolutionary divergence from other eukaryotes, likely driven by selective pressures encountered in its parasitic lifecycle within mammalian hosts and tsetse fly vectors. This divergence highlights how *T. brucei*’s axoneme evolved to optimize motility for infection and transmission. Despite its central importance to axoneme structure and motility, however, an atomic description of *T. brucei*’s central apparatus remains unknown.

In this study, we utilized a combination of single particle cryoEM, mass spectrometry analysis, and Alphafold3-based modeling to examine the structure of the *T. brucei* CA. Our structure revealed seven conserved protein densities, among which PF16 emerges as a key architectural element. While the outer doublet microtubules are arranged with radial symmetry with the canonical 9+2 pattern, asymmetry arises from the non-identical projections and protein composition of the central apparatus microtubules (C1 and C2). This asymmetry is fundamental for generating *T. brucei*’s distinctive bihelical waveform (20) and directional motility. Our findings highlight PF16’s role in anchoring and propagating this asymmetric organization across the CA. Notably, we also characterize the intrinsic molecular flexibility of the central apparatus, previously proposed in other systems, now apparent in the different conformations of PF16. In addition, we identified *T. brucei*-specific densities, suggesting species-specific adaptations linked to the parasite’s unique flagellar dynamics. These findings deepen our understanding of *T. brucei* flagellar architecture and highlight PF16 as a potential mediator of axoneme dynamics and contributor to the 9+2 axoneme asymmetry.

## Results

### Structure determination of *T. brucei* central apparatus

Previously, we developed a workflow to split the *T. brucei* flagella into individual DMTs for cryoEM analysis (18). The dataset used to resolve the split DMT structure also contained clear images of the central apparatus (Figure S1A-B). Taking advantage of this, we further analyzed this dataset to reconstruct the C1 and C2 singlet microtubules using single-particle cryoEM. We manually screened and picked images with singlet microtubules: from the 8425 micrographs, 526140 particles were selected for further analysis. Following an initial global angular search, 3D classification was used to distinguish C1 and C2 microtubules, resulting in global reconstructions at ∼3.8 Å for C1 and ∼4.4 Å for C2 16-nm repeats, and further classification did not reveal 32-nm periodicity in either the C1 or C2 microtubules (Figure S1C). Due to the flexible nature of C1 and C2 components, it was challenging to resolve proteins located further away from the tubulin lattice. To overcome this limitation, we employed a combination of center shifting, tubulin density subtraction, and local refinement to better resolve peripheral regions of the CA (Figure S1C). These strategies allowed us to obtain a composite map that revealed detailed structural features of these otherwise elusive components (Figure 1A, 1B).

**Figure 1.**
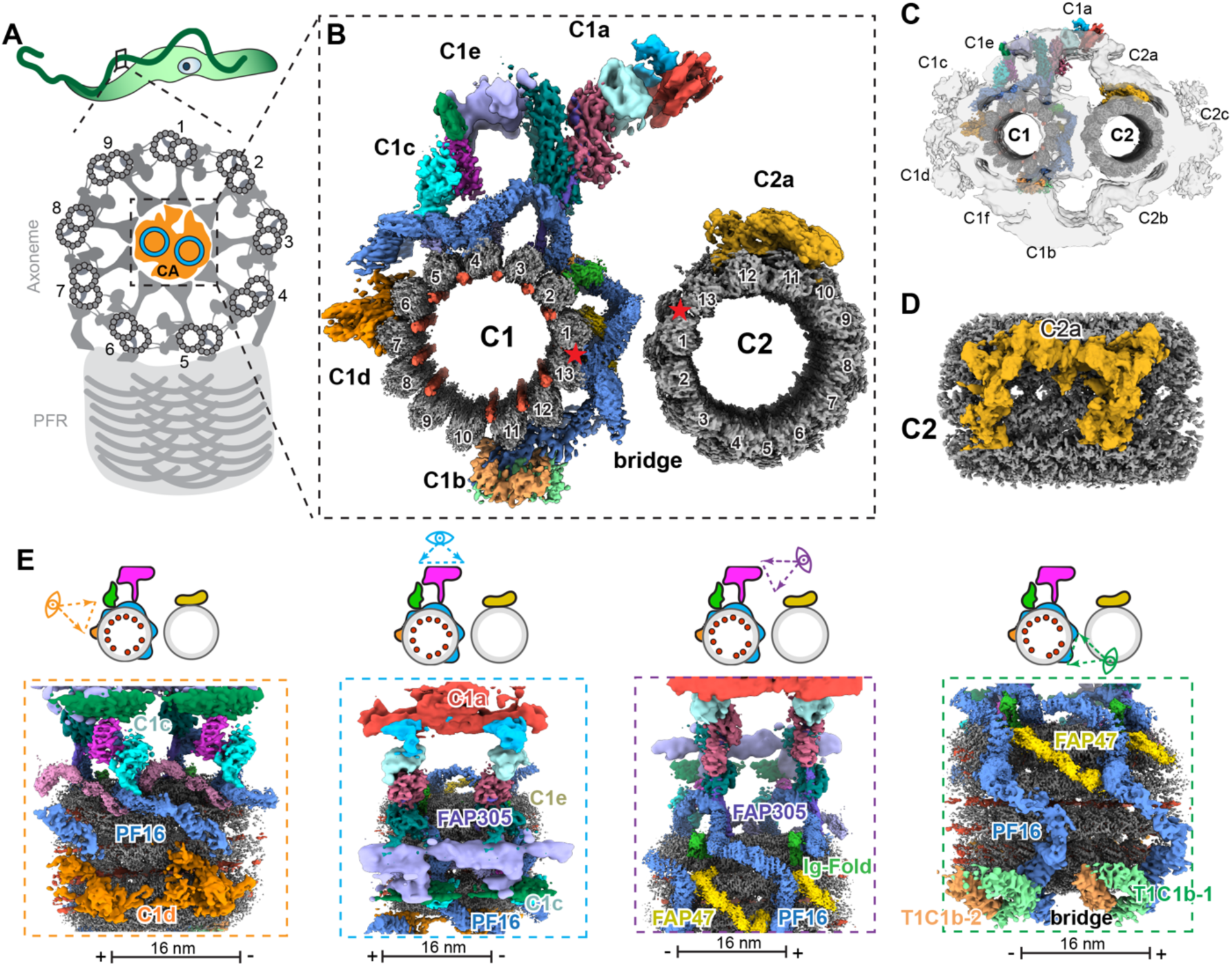
Cryo-EM reconstruction and regional segmentation of the *Trypanosoma brucei* central apparatus. **(A)** Schematic of the flagellum from *T. brucei*, indicating the central apparatus (CA), peripheral doublets, and the paraflagellar rod (PFR). **(B)** Composite map of cryoEM reconstruction of C1 and C2 microtubules. Positions of microtubule seams are indicated by red stars. **(C)** Superposition of 3D reconstructions of C1 and C2 microtubules with low resolution cryoET map of *T. brucei*. Projections of C1 and C2 are labeled. **(D)** C2 microtubule with resolved density for the C2a projection. **(E)** Labeled CA subregions, including C1a, C1c, C1d, C1e, and bridge domains, revealing positional organization across the C1–C2 interface.

To validate the overall architecture of the CA, we docked our high-resolution structure into a previously published lower-resolution cryoET map (16) (Figure 1C). This confirmed that each protein density occupied its expected location within the intact axoneme. Among the C1 projections previously established, prominent densities corresponding to the C1a, C1c, and C1e projections are clearly present, whereas most components of the C1b, C1d, and C1f projections are not observed (21). For the C2 microtubule, the C2a arm is resolved, but densities corresponding to C2b and C2c were absent (Figure 1D). The missing densities are likely due to our sample preparation workflow, which involves protease digestion to separate DMTs and may have removed certain proteins of the CA. We took advantage of this simplified structure to identify the most strongly associated components of the CA assembly and organization. Consistent with prior tomography studies in *T. brucei*, the CA proteins identified by cryoEM exhibit a 16-nm periodicity, matching the continuous helical wrapping of PF16 around the C1 microtubule. Our high-resolution maps enabled accurate atomic modeling of tubulin and PF16. For proteins with lower-resolution density, we fitted AlphaFold-predicted homologs (22) based on previously characterized CA structures in *Chlamydomonas* (*8, 9*). In cases where no clear homologs were available, we employed DomainFit (23) to semi-automatically screen candidate proteins from a published *T. brucei* flagella mass spectrometry dataset (18), selecting the most likely matches based on their fit scores to the unassigned densities. Altogether, 3 proteins on C1 (PF16, FAP47, FAP305) and 1 protein on C2 (FAP65) were confidently assigned to the cryoEM densities, along with α- and β-tubulin, which form the lattice structure of both C1 and C2 microtubules (Figure 1E). Of the Tb-specific proteins in our structure, we were able to assign potential candidates to three densities using DomainFit, which we have termed TbC1b-1 (candidate Q57WS7), Tb-C1b-2 (Q38HH0/Q4GYV5), and Ig-fold like protein (Q385G3, Q388P3, Q57XV2, Q388P3, Q383L3) (Supplementary Figure 2).

Although PF16 is structurally conserved and prominently decorates the C1 microtubule, many other densities differ significantly from those observed in the central apparatus of *Chlamydomonas* (*8, 9*). The *Chlamydomonas* CA contains three conserved armadillo repeat proteins: FAP194, FAP69, and PF16. In *T. brucei*, however, FAP194 and FAP69 are non-homologous to any flagellar proteins and instead, distinct helical densities, TbC1b-1 and TbC1b-2, occupy the similar locations on C1 (Figure 1E). Additionally, within the C1c region, the architecture, similar to that seen in proteins like Fap266 and Fap227.

Within the C1 microtubule lumen, we observed 11 filamentous structures decorating the furrows between adjacent tubulin protofilaments (pf) (Figure 1B). These are found between all pfs, with the exception of pf1/13 and pf9/10. These filamentous MIPs bind to the microtubule lumen in a similar location to RIB43, which stabilizes protofilaments in the pfA11 to pfA13 “ribbon” region of doublet microtubules (24) (25). Notably, we did not observe well-defined globular densities within either C1 or C2, consistent with the *Chlamydomonas* C1 structure, which contains only microtubule inner protein (MIP) densities (9). In contrast, this differs from the *Chlamydomonas* C2 structure, which includes prominent globular proteins such as FAP213 and FAP196. This absence in *T. brucei* may reflect a true lack of large globular domains in its central microtubules or, alternatively, the loss of such domains during our sample preparation.

### Spiral arrangement and structural flexibility of PF16 support central apparatus organization

PF16 is a key organizer of the C1 microtubule and plays a critical role in maintaining the relative orientation between the C1 and C2 microtubules. In *T. brucei*, PF16 forms an organized network of segmented, right-handed helices that coil along the cylindrical surface of the microtubule, resembling a vine spiraling around a pole (Figure 2A-C). This arrangement forms distinctive spirals in our structure spanning from the C1d region, wrapping around the microtubule in a clockwise manner, looking toward the flagellum tip, toward the C1b region, where it ends near the two distinct helical densities Tb-C1b1 and Tb-C1b2 (Figures 1E, 2B, 2C). Unlike *Chlamydomonas*, which contains two interleaved PF16 spiral types (type I and II) that form a discontinuous, triple-helical pattern capped by FAP194 (8, 9), *T. brucei* displays a single, continuous spiral that repeats every 16 nm along the C1 microtubule. In this way, *T.brucei* shows PF16-specific differences in CA projection architecture and periodicity.

**Figure 2.**
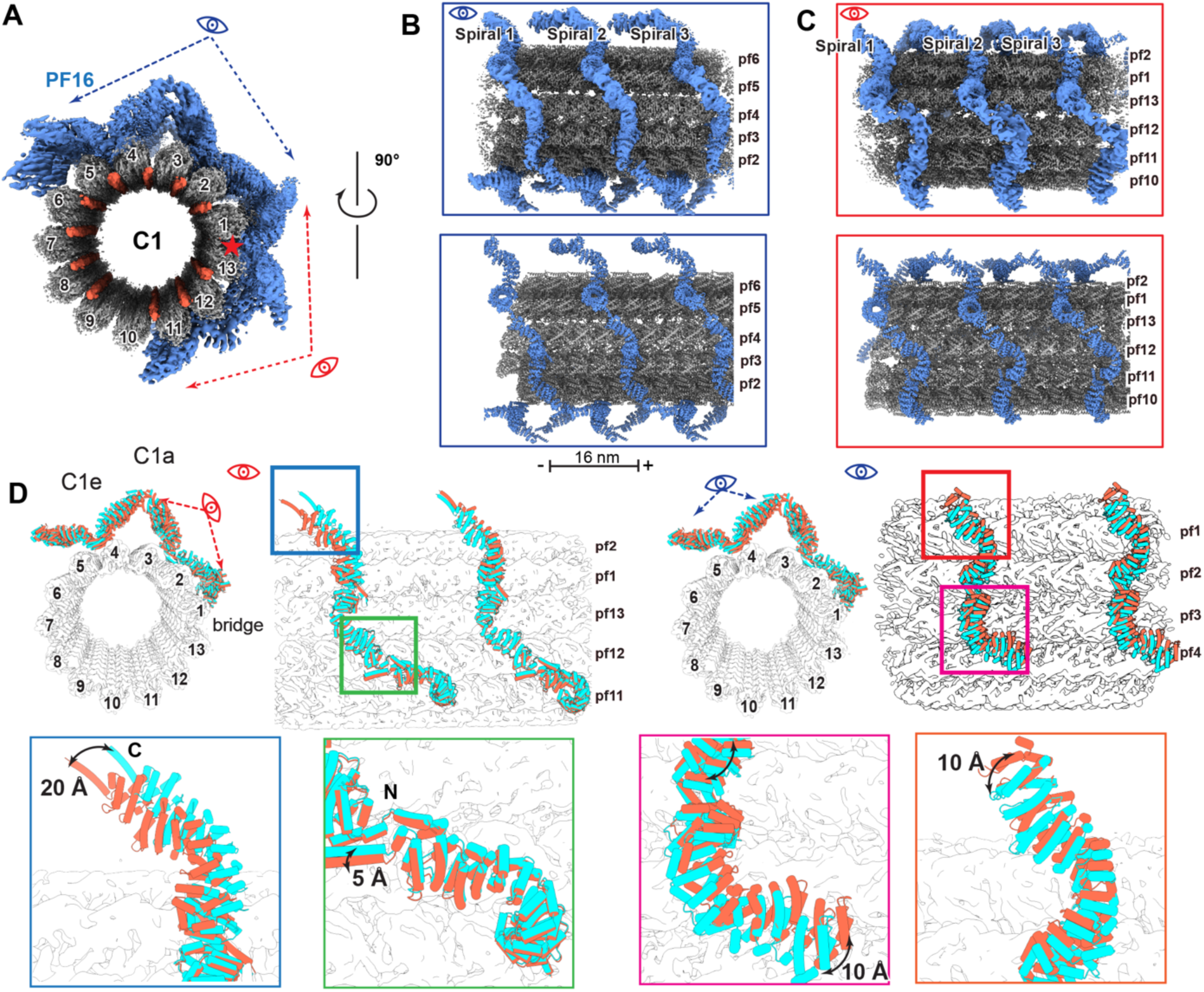
PF16 forms a flexible helical scaffold that spans the surface of the C1 microtubule. **(A)** PF16 (blue) wraps around the surface of the C1 microtubule (tubulin in gray). **(B-C)** CryoEM map (upper panel) and atomic model (lower panel) of PF16 spiral viewed from different angle as indicated in (A). **(D)** PF16 adopts different conformations depending on its position along the C1 microtubule, demonstrating its elastic adaptation to curvature.

Armadillo repeat (ARM) proteins like PF16 are structurally elastic, allowing substantial changes in tertiary conformation while the secondary structure remains relatively unchanged (26). Such elastic capacity is relevant for CA structures, which must accommodate extreme and dynamic structural changes and dynamic interactions with radial spoke heads as the structure bends and twists during axoneme beating. Previous work in *Chlamydomonas* compared 20 PF16 subunits within a single CA repeat, revealing large tertiary structural variations, particularly in the relative positions and orientations of PF16 monomers bound to different protofilaments (8). However, it remained unclear how PF16’s secondary structure adapts to accommodate these tertiary structural shifts, to what extent PF16 undergoes conformational changes across the microtubule surface, and whether specific regions within the PF16 spirals are more structurally dynamic than others.

To address these questions, we performed 3DFlexibility Analysis (27) across the well-resolved PF16 regions in the C1c, C1e, C1a, and bridge regions (Figure 2D). The most pronounced conformational changes were observed in PF16 monomers in the C1a arm, with displacements reaching up to 20 Å, highlighting significant flexibility in this projection-rich region. In contrast, PF16 within the bridge region displayed minimal positional deviation, consistently ranging between 2 and 4 Å, whereas intermediate levels of flexibility were observed in the C1e and C1c regions, where PF16 exhibited shifts of up to 10 Å. Consistent across the C1 microtubule, the C-terminal ends of PF16, which polymerize above the microtubule and are involved in recruiting central apparatus projection proteins, showed greater conformational variability. In contrast, the ARM repeats near the tubulin interface and the N-terminal dimerization regions remained relatively rigid. This spatial pattern of flexibility suggests that PF16 maintains a tight association with the microtubule lattice while permitting movement at distal ends to accommodate dynamic structural changes during axonemal beating and shifting interactions between CA projections and the radial spokes.

### PF16 hydrophobic dimerization domains and tubulin electrostatic interactions promote stability and assembly

Given the substantial tertiary structural shifts that PF16 accommodates across the CA, its stable incorporation into the microtubule scaffold must rely on strong, conserved interfaces (Figure 3A). Dimerization of neighboring PF16 monomers occurs through their N-terminal tails, which adopt an antiparallel configuration—residues 1–20 of one subunit align and interact with residues 20–1 of the opposing subunit (Figure 3B). This antiparallel arrangement results in a palindromic hydrophobic interface enriched in aromatic residues (Figure 3C). A prominent feature of this interface is the reciprocal interactions between aromatic side chains of Y13 and F20 on opposing strands. These residues engage in parallel, face-to-face interactions, where the planar aromatic rings stack tightly and contribute to a repeating FFYYFF motif that defines the hydrophobic core of the N-terminal interface. Additionally, hydrophobic residues such as I6, V9, V24, and F27 form complementary stacking interactions on either side of the central π–π stacking interface, further reinforcing the hydrophobic core and stabilizing the PF16 dimer (Figure 3B, 3C). This highly ordered, interdigitated arrangement promotes the structural rigidity of the PF16 dimer where it contacts the tubulin lattice, reinforcing its role as a stable structural foundation within the C1 microtubule.

**Figure 3.**
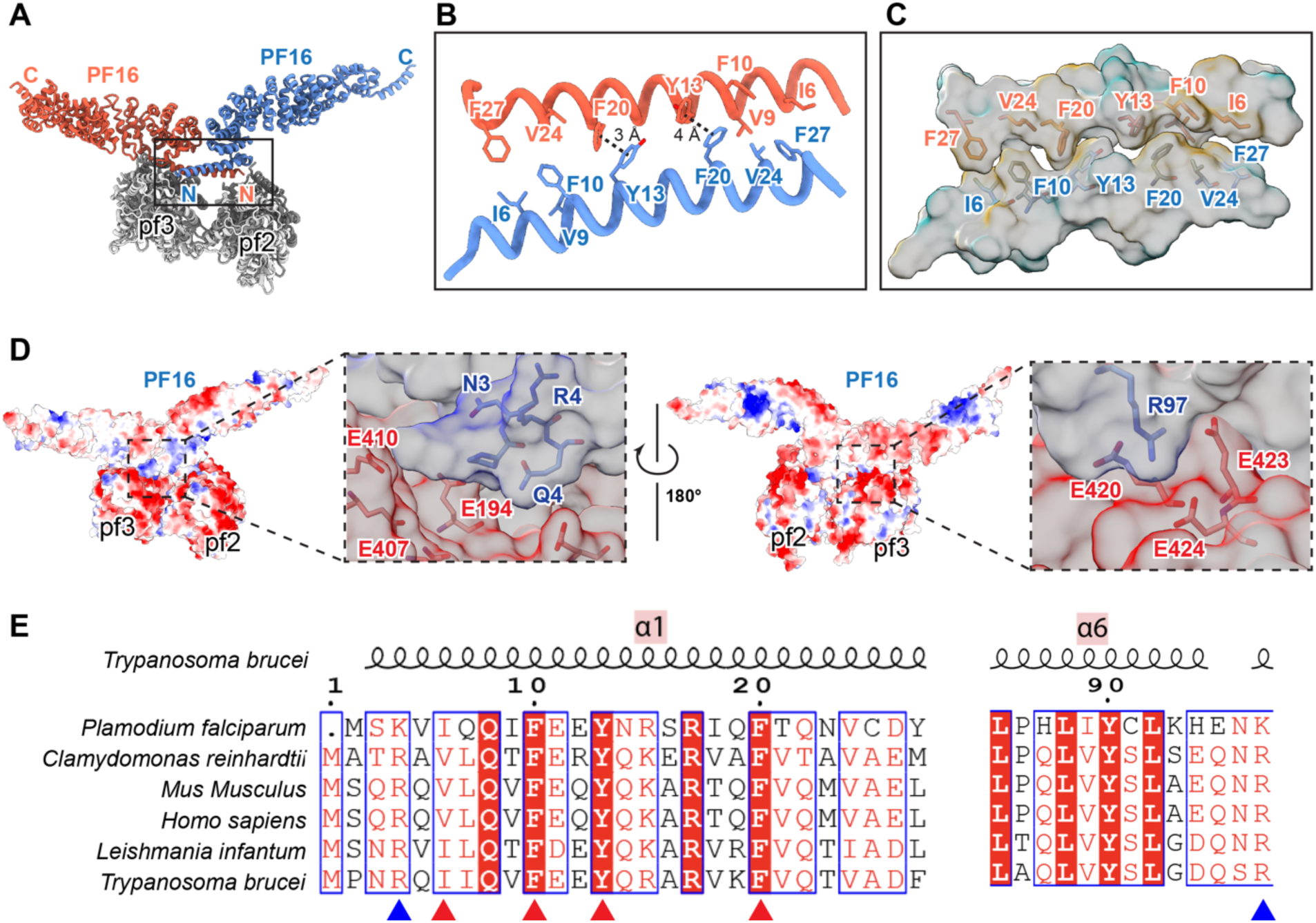
N-terminal dimerization of PF16 mediates inter-subunit stabilization and microtubule interaction. **(A)** PF16 dimer shown with one monomer in red and the other in blue, illustrating their antiparallel helical interface. **(B)** Zoomed view of the N-terminal dimerization interface, highlighting tight interdigitation of hydrophobic residues. **(C)** Hydrophobic interaction network stabilizing the dimer, including residues I6, V9, Y13, V24, F27. **(D)** Electrostatic surface representation of PF16-tubulin interaction interface reveals charged patches aligned with tubulin’s surface. **(E)** Sequence alignment of PF16 orthologs from trypanosomes, humans, and algae reveals strong conservation at the dimer interface and tubulin-binding surface.

Within and around the region of dimerization, PF16 also engages in extensive electrostatic interactions with the underlying tubulin lattice. Positively charged residues in PF16 interact with negatively charged, glutamate-rich patches on the tubulin surface (Figure 3D). At the very N-terminus of PF16, R4 binds within an acidic groove formed by residues near the E410 region of tubulin, while R97, located on a later helix of PF16, interacts with another glutamate-rich region around E420-E424 of tubulin. These interactions position PF16 in precise alignment along the protofilament and contribute to its robust attachment to the microtubule wall. Together, the hydrophobic interactions that stabilize the PF16 dimer and the electrostatic interactions that mediate its attachment to tubulin cooperatively ensure the stable incorporation of PF16 into the central apparatus. Supporting the fundamentally important role of these interactions, sequence alignment of PF16 across a broad range of protist and metazoan species reveals strong conservation of key residues—including R4, F10, Y13, F20, and R97— underscoring their evolutionary importance in maintaining CA structure and function (Figure 3E).

### Structure comparison reveals conserved proteins and lineage-specific adaptations

Previous studies of CA structure in *Chlamydomonas* revealed proteins with diverse periodicities, typically aligned to the 8-nm repeat of the tubulin lattice, with the longest periodicity reaching 32 nm (8, 9). In contrast, our structural analysis of the *T. brucei* CA reveals a strikingly consistent 16-nm periodicity, with densities arranged atop the PF16 spirals that wrap around the C1 microtubule. By combining information from proteins resolved in our cryoEM structure and proteins identified via mass spectrometry (18), we found 28 CA proteins conserved between *T. brucei* and *Chlamydomonas*. Among the 26 conserved proteins between *Chlamydomonas reinhardtii* and *Trypanosoma brucei* (excluding tubulin and PF16), 4 localize to the C1a projection (FAP101, Calmodulin, FAP305, FAP119), 5 to C1e (FAP81, FAP15, FAP119, FAP227, PF6), 2 to C1c (FAP81, FAP65), and 3 to C1f (FAP266/CARP10, FAP74, FAP114). For the C1d region, 4 conserved proteins were identified (FAP15, FAP42, FAP246, PF6), and 3 proteins are shared in the C1b projection (Hydin, CPC1, Enolase [dimer]). Within the C2 microtubule, 5 conserved proteins were found in C2a/e (FAP70, FAP279, FAP20, FAP65, and PF20) (Table S1; Figure 4A–C). This analysis highlights the C1a, C1c, and C1e subregions as the most conserved, underscoring their likely critical roles in CA structural organization and function.

**Figure 4.**
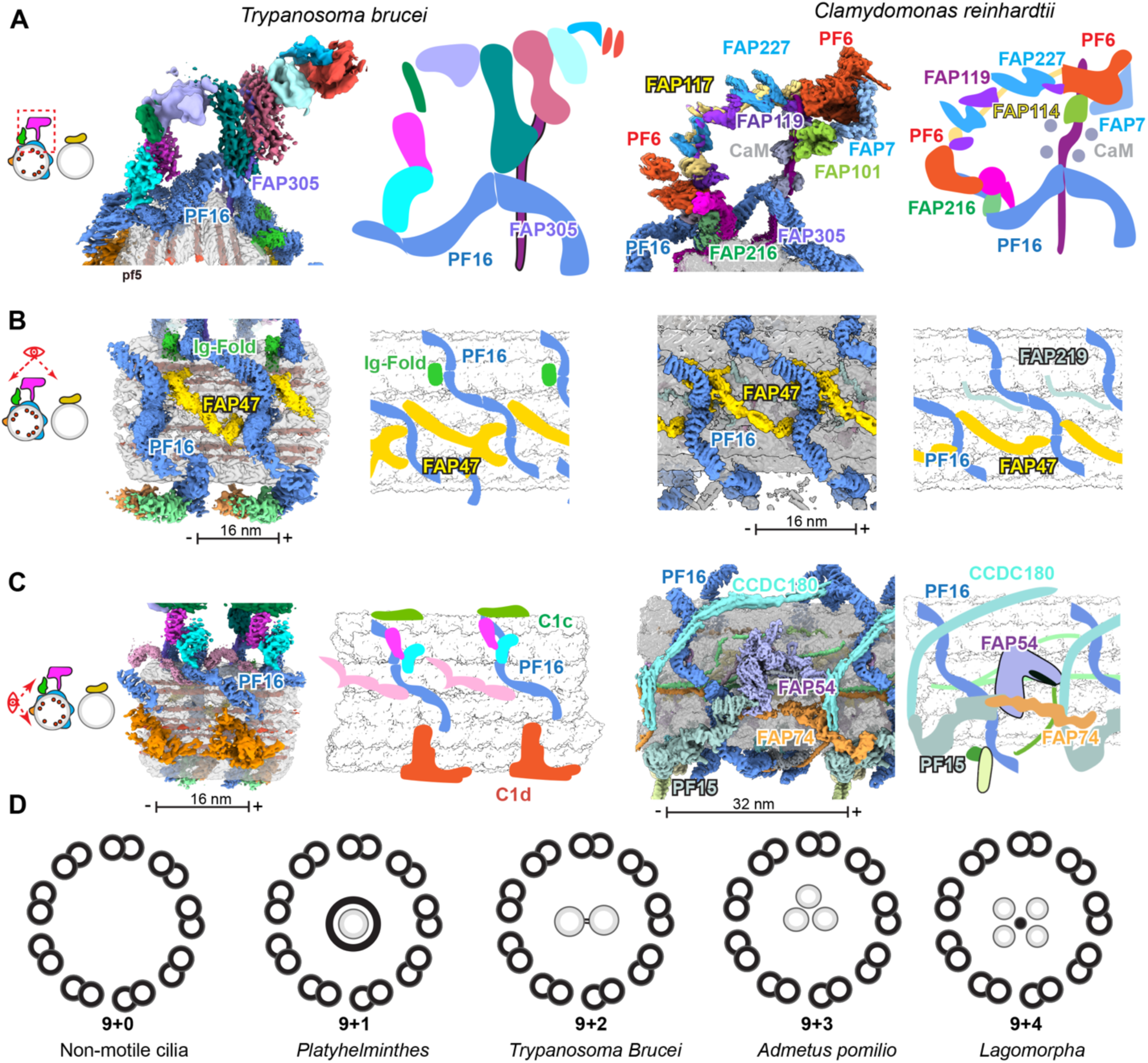
Comparative architecture of central apparatus projections between *T. brucei* and *Chlamydomonas*. **(A)** Structural comparison of the C1a and C1e projections in *T. brucei* and *Chlamydomonas reinhardtii* reveals species-specific protein incorporation and domain organization. **(B)** Differences in the bridge region morphology and PF16–FAP305–FAP47 arrangements between species, highlighting evolutionary tuning of CA connectivity. **(C)** Schematic comparison of C1c and C1d projections between *T. brucei* and *C. reinhardtii*, showing distinct spatial arrangements. (**D**) Schematic representation of axoneme organization from different organisms, showing the “9+n” architecture.

Conversely, several *Chlamydomonas* proteins—FAP7, FAP46, FAP54, FAP69, FAP194, FAP105, FAP108, FAP213, FAP236, FAP219, FAP216, FAP147, FAP174, FAP221, FAP239, FAP275, FAP388, FAP297, FAP225, FAP424, and HSP70A—have no clear homologs in *T. brucei*, suggesting lineage-specific protein loss or replacement. The C1 b, d, and f regions, in particular, shows the significant divergence, marked by the absence of corresponding densities and the presence of non-homologous proteins (28) (Figure 4C). Most notably, the composition of microtubule inner proteins (MIPs) varies dramatically: *Chlamydomonas* contains arc-MIPs and SAXO proteins, such as FAP105, FAP236, and FAP275, that form lateral arcs or longitudinal arrays within the lumen, often stabilizing specific protofilament interfaces and remodeling tubulin loops (9). In contrast, *T. brucei* lacks these MIPs in the genome and instead features a distinct set of longitudinal, rod-like MIPs resembling RIB43a, which align tightly along protofilaments and reflect a fundamentally different strategy of microtubule stabilization and inner scaffold architecture.

Structurally, many conserved CA proteins share recurring domain architectures that hint at common mechanistic roles. FAP81, FAP47, and FAP65 contain immunoglobulin (Ig)-like folds, β-sandwich motifs that provide both mechanical stability and flexibility across protofilament interfaces (29). PF16 consists of multiple armadillo repeats that form a helical solenoid capable of conforming to microtubule curvature while maintaining rigidity. Calmodulin, widely distributed within the CA, features EF-hand motifs that bind calcium and undergo conformational shifts, serving as molecular switches for motility regulation. Additional conserved proteins such as FAP101, FAP15, and FAP266 include HEAT or coiled-coil domains, which facilitate protein–protein interactions and contribute to the flexible linkage between distinct CA substructures. (30) (31). The recurrence of these motifs suggests that the CA is a modular structure composed of conserved scaffolding and signaling components that have been selectively tuned across species to accommodate diverse modes of flagellar beating.

Interestingly, while the C1a and C1e regions exhibit structural divergence between *T. brucei* and *Chlamydomonas*, the C1 bridge is remarkably conserved (Figure 4B). This bridge is built from 16-nm PF16 repeating spirals interspaced by FAP47 monomers, forming a regular architectural unit. Despite subtle organizational differences, this arrangement is preserved between *Chlamydomonas* and *T. brucei*. Together, these demonstrate that while the *T. brucei* CA retains the conserved modular PF16 scaffold, it has diverged structurally through presence of species-specific MIPs and the replacement or absence of *Chlamydomonas*-specific proteins, adaptations that likely support its unique helical motility.

## Discussion

The current study resolved the structure of the CA from *T. brucei*, a eukaryotic pathogen featuring a non-rotating CA within a 9+2 axonemal configuration. The cryoEM structure reveals a highly organized CA architecture comprised of periodic PF16 spirals as a foundation for asymmetrically distributed bridge proteins. Through structural mapping and protein identification, we establish PF16 as a central organizing hub that imparts both stability and asymmetry to the CA, while also enabling structural flexibility required for effective flagellar beating. These insights provide a mechanistic framework for understanding how the CA maintains orientation and regulates motility in non-rotating axonemal systems.

Our structural analysis shows that PF16 possesses spring-like tertiary flexibility near its C-terminal domains while remaining rigid in its N-terminal regions. We define amino acids responsible for rigidity of N-terminal overlaps in adjacent PF16 molecules as well as those responsible for maintaining contact with tubulin of pfs. PF16’s combined flexibility plus stable attachment to the microtubule, allows the CA to absorb mechanical stresses and adapt to dynamic changes in the flagellum without losing its alignment. Flexibility of PF16 may enable the CA to maintain a consistent orientation relative to the outer doublets ensuring that mechanical and regulatory signals are correctly communicated to the different dynein arms across the axoneme. Indeed, loss of PF16 in causes the CA to rotate within the DMTs with concomitant erratic flagellar beating (32). The importance of PF16 in CA architecture is further supported by the distribution of proteins that associate with it: the C1a, C1c, and C1e subregions, which are associated with PF16 density, are the most structurally preserved in our structure and homologous across species. In contrast, C1d and C1f, lacking clear PF16 support in our map, are notably more divergent and mainly absent in our structure, suggesting that PF16 contributes to structural conservation in the CA and may be needed to facilitate recruitment of associated CA proteins.

Functional studies in *T. brucei* and other organisms reinforce PF16’s role in CA assembly and function. In *Chlamydomonas reinhardtii*, *pf16* mutants exhibit severe flagellar defects, with ∼60% of axonemes lacking the CA entirely (9+0), ∼20% showing a 9+1 arrangement, and only ∼20% retaining the typical 9+2 configuration (33) (34). Similarly, in *Plasmodium berghei*, PF16 disruption results in the complete loss or partial assembly of the central apparatus, indicating a critical role in initiating or stabilizing central apparatus formation (35). Strikingly, in *T. brucei*, PF16 loss does not prevent CA assembly, as RNAi knockdown of PF16 leads to predominantly intact 9+2 structures, but it does cause misalignment of CA microtubules, resulting in loss of their fixed orientation (32). These findings are consistent with reports in mammalian sperm, where PF16 contributes more to CA orientation than structural integrity (36).

Taken together, these studies reveal a unifying theme: PF16 is essential for introducing and maintaining the asymmetric geometry of the central apparatus that underpins regulated dynein activity and waveform propagation. In organisms like *Chlamydomonas* and *Plasmodium*, which either rotate their CA or rely on dynamic flagellar beating for rapid gamete release, PF16 is critical for the physical assembly and stability of the central apparatus. In contrast, in *T. brucei* and mammalian sperm—organisms with a fixed CA orientation—PF16 is not strictly required for assembly but rather for maintaining orientation and coordinated motion, with its loss leading to disorganized and inefficient motility. This dichotomy underscores PF16’s dual role: as a structural stabilizer of the 9+2 axonemal symmetry and as a spatial orienter of the central apparatus in non-rotational systems. Our work suggests that in *T. brucei*, PF16 and associated proteins, such as FAP47, introduce and stabilize the C1–C2 asymmetry by bridging between the two microtubules at regular 16-nm intervals. We propose that PF16 serves as a nucleation point for the bridges connecting the CA, anchoring C1 and guiding the periodic placement of C2- linked components. The loss of CA asymmetry due to PF16 deficiency disrupts the CP-radial spoke regulatory axis responsible for spatial regulation of dynein motors around the axoneme (37), leading to uncoordinated or ineffective flagellar movement.

The architectural asymmetry of the central apparatus is a fundamental determinant of axonemal function (37). In typical 9+2 systems, the two central microtubules define a bilateral plane that coordinates radial spoke–dynein interactions and waveform propagation (38). In systems with non-canonical arrangements (e.g., 9+3 in crustaceans and insects, 9+4 in rabbit embryos), altered central symmetry reflects unique functional adaptations (39) (40) (41). The number of central microtubule filaments—0, 1, 2, or 3—within the axoneme significantly impacts its structural symmetry, which in turn influences its functional properties. In the canonical, 9+2 axoneme architecture, the two central microtubules establish a plane of bilateral symmetry, dividing the nine outer microtubule doublets into two halves and facilitating coordinated dynein activity for symmetrical, wave-like ciliary beating (Figure 4D) (38). This symmetry is disrupted in 9+0 configurations, where the absence of a CA, as in non-motile primary cilia, eliminates a defined central axis, often correlating with a lack of rotational or planar motion and a more radially symmetric structure suited for sensory roles. Axonemes with one (9+1) or three (9+3) central microtubules, seen in certain mutants or species like some flatworms and arachnids (42) (43), introduce asymmetry or an uneven central framework, which can skew the radial spoke interactions and lead to irregular or less efficient beating patterns. Thus, the central microtubule count directly shapes the axoneme’s symmetry, balancing structural stability and dynamic function according to the cilium’s biological role. These deviations from the 9+2 pattern highlight the diversity of axonemal structures in nature, particularly in arthropods, where evolutionary adaptations have led to specialized sperm morphology and other cilia.

In conclusion, our study provides the most detailed view to date of the *T. brucei* CA. We demonstrate that PF16 is a key architectural determinant that mediates assembly, asymmetry, and flexibility of the CA and define amino acids responsible for PF16 dimer contacts and microtubule attachment. These features are essential for the spatial regulation of motility in non-rotating cilia. By characterizing lineage-specific and conserved CA densities, we uncovered broader principles governing axonemal organization and suggest new avenues for targeting flagellar function in pathogenic protists.

## Materials and Methods

### CryoEM grid preparation and data collection

Procyclic (insect midgut stage) *Trypanosoma brucei brucei* (strain 29-13), originally obtained from George Cross (Rockefeller University) (44), were cultured in SM medium (45) supplemented with 10% heat-inactivated fetal bovine serum (FBS) at 28°C in a humidified incubator with 5% CO_2_. CryoEM analysis was performed using the same dataset and cryoEM grids described in our previous publication (18). Briefly, demembranated flagellar skeletons were isolated and resuspended in Splitting Buffer (20 mM HEPES, pH 7.4, 3 mM MgCl_2_, 0.1 mM EGTA, 1 mM DTT, 25 mM KCl) without protease inhibitors. The sample was mildly digested with 10 µg/mL trypsin (Sigma-Aldrich, T4799) on ice for 20 minutes. Proteolysis was halted by the addition of 2× SigmaFAST EDTA-free protease inhibitor cocktail (Sigma-Aldrich, S8830). The sample was then sonicated for 30 seconds at 4°C in a bath ultrasonic cleaner (BRANSON), fragmenting the long flagella into shorter segments of approximately 2–3 µm. The fragmented sample was pelleted and resuspended in 40 µL of Splitting Buffer supplemented with protease inhibitors. To dissociate individual doublet microtubules (DMTs), 2.5 mM ATP was added, and the sample was incubated at room temperature for 25 min. The split sample contained all components of *T. brucei* flagella, including DMTs, CA, and PFR. Freshly prepared samples were immediately used for cryoEM grid preparation.

To prepare cryoEM grids, 3 µL of the DMT-containing solution was applied to a glow-discharged holey carbon grid (Quantifoil 2/1; Electron Microscopy Sciences). After a 30-second incubation, the grid was blotted for 9 seconds at 4°C with a blot force of 4 and 100% humidity using a Vitrobot Mark IV (Thermo Fisher Scientific). The grid was then plunge-frozen in liquid ethane and stored in liquid nitrogen until imaging. Data were collected on a Titan Krios transmission electron microscope (Thermo Fisher Scientific) operating at 300 kV. Images were recorded at a nominal magnification of 81,000× in super-resolution mode, corresponding to a calibrated pixel size of 0.55 Å at the specimen level. A Gatan energy filter with a slit width of 20 eV was used. Movies were acquired using SerialEM 4.0 (46) with an exposure time of 2 seconds and a total electron dose of ∼45 electrons/Å², dose-fractionated over 40 frames. The defocus range was set between −1.6 and −2.4 µm. The 40 frames within each movie were aligned in MotionCor2 (47) with the first and last frame excluded. The images were binned 2× to a pixel size of 1.1 Å. The defocus values of micrographs were determined by CTFFIND4 (48). Of the total of 106,694 movies collected for DMT structure determination, 8425 images were manually screened and picked for CA reconstruction.

### CryoEM image processing and 3-dimensional reconstruction

Particles were picked along the microtubule by Topaz (filament tracing) (49, 50) and extracted by an 8-nm interval. After 2D classification, we initially performed a global refinement with large search range parameters defined in the MIRP protocol (51) to define basic angular and translational assignments. We did a broad global search to assign appropriate rotational angular assignments before a round of classification to separate the singlets into C1, C2, and singlet microtubules. We then performed initial averaging and alignment in Relion (52) before importing the data into cryoSPARC (53) for further local refinement. We employed a comprehensive strategy of recentering particles, performing local refinement, and particle subtraction to ideally refine outer densities in several areas. We combined several maps that had been locally refined to create our global map. The resultant cryo-EM maps of the 16 nm repeating C1 Central Apparatus exhibited a global resolution of 3.8 Å with local resolutions ranging from 3.5-3.8 Å and C2 with a global resolution of 4.4 Å with local resolutions ranging from 4.0 to 4.4 Å. We then used Alphafold3 to fit potential protein candidates. We also collected mass spectrometry data from purified flagellum skeletons to produce a library of potential *T*. *brucei* flagellum proteins (18) and utilized cryoID to identify most likely candidates for certain map densities (54). AlphaFold-predicted structures served as initial models for atomic modeling of both conserved and species-specific cryoEM map densities(55, 56)

### Atomic modeling and docking

The tubulin models were built using AlphaFold-predicted models of α- and β-tubulin and using molecular dynamics flexible fitting software in UCSF ChimeraX (57, 58). We distinguished α- and β-tubulin based on a conserved loop structure on the inner side of the microtubule singlet which is shorter in β-tubulin (59). This distinct structural difference is readily identifiable in our map and aided in locating the C1 seam. The identities of unknown densities were confirmed using automated building in ModelAngelo and standard Protein BLAST of the predicted amino acid sequences against the TriTrypDB database(60–62). Alternatively, or often in combination with ModelAngelo-predicted models, cryoID was used to identify the most likely candidates for cryo-EM densities(54). Further attempts to fit proteins in low resolution regions were made using a strategy similar to that of the DomainFit software package(63). Briefly, visual inspection of AlphaFold-predicted structures also aided in matching of candidates with map density shapes to assess potential matches. Models were fit using Coot and ISOLDE as described previously (57, 64) and refined using Phenix Real Space Refinement (65).

## Data Availability

The cryoEM density maps were deposited in the Electron Microscopy Data Bank (EMDB) with accession code EMDB-XXXX. The corresponding models were deposited in the Protein Data Bank (PDB) with code XXXX. This paper does not report original code. All other data needed to evaluate the conclusions are present in the paper and/or the supplementary materials.

## Acknowledgements

This project is supported by grants from the US NIH (R01GM071940 to Z.H.Z.; R01AI052348 to K.L.H.). We acknowledge the use of resources in the Electron Imaging Center for Nanomachines supported by UCLA and grants from the NIH (1S10OD018111).

## Author Contributions Statement

Z.H.Z. and K.L.H. conceived the project and supervised research; X.X. prepared samples, recorded cryo-EM images, S.K. and X.X. processed the data and analyzed the structure; S.K. and X.X. interpreted the results and wrote the manuscript; and all authors reviewed and approved the submitted paper.

## Supplementary Figures

**Supplementary Figure 1.**
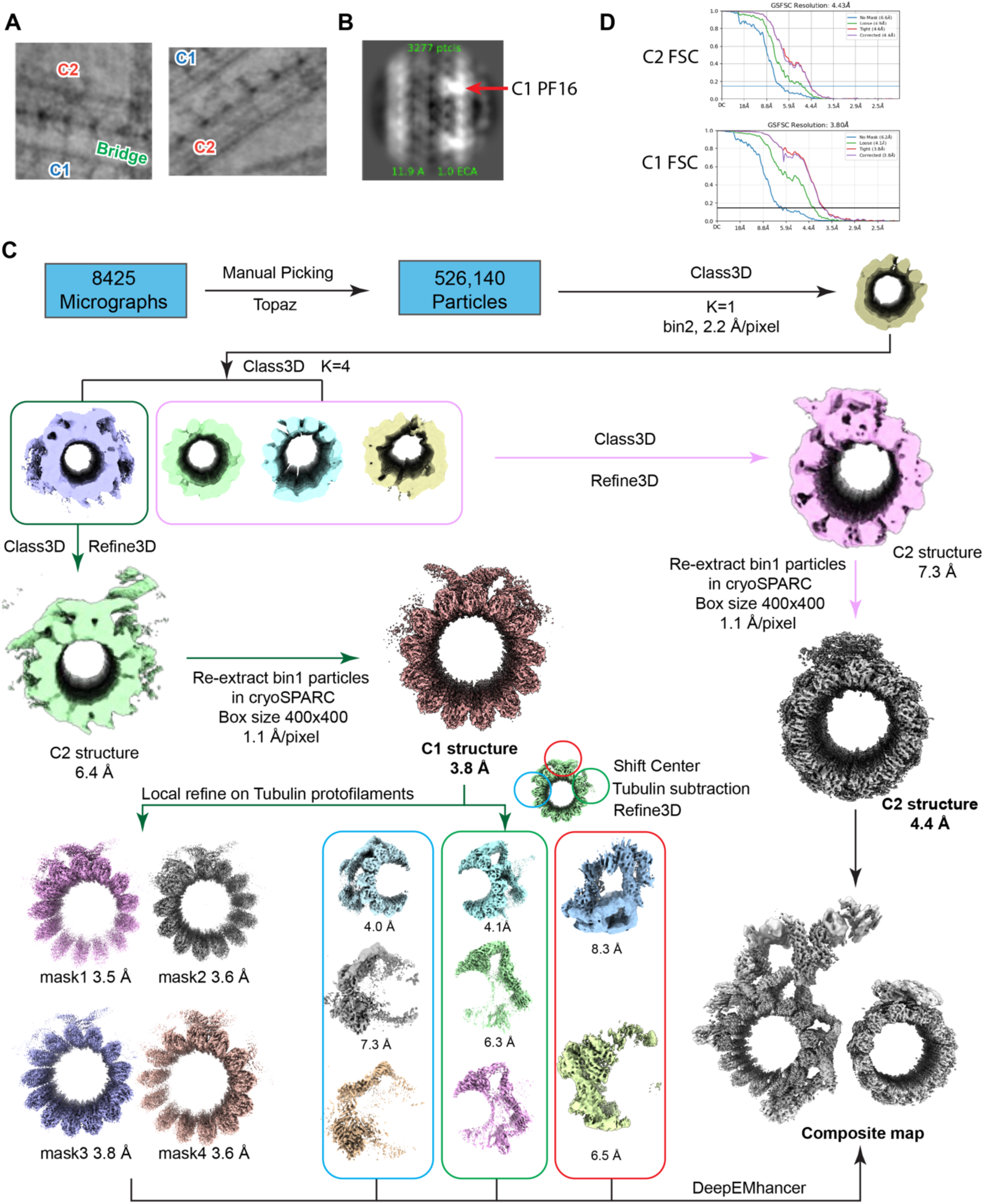
Cryo-EM processing workflow for resolving *T. brucei* central apparatus microtubules. (**A**) Representative cryoEM images of central pair. (**B**) One image from 2D classification showing the density of PF16. (**C**) Stepwise strategy for particle picking, 2D and 3D classification, and local refinement to achieve high-resolution reconstructions of C1 and C2 microtubules. Final reconstructions include maps for CA-tubulin interfaces and bridge components at resolutions ranging from 3.5–8.3 Å. (**D**) Fourier Shell Correlation (FSC) curves of C1 and C2 reconstruction.

**Supplementary Figure 2.**
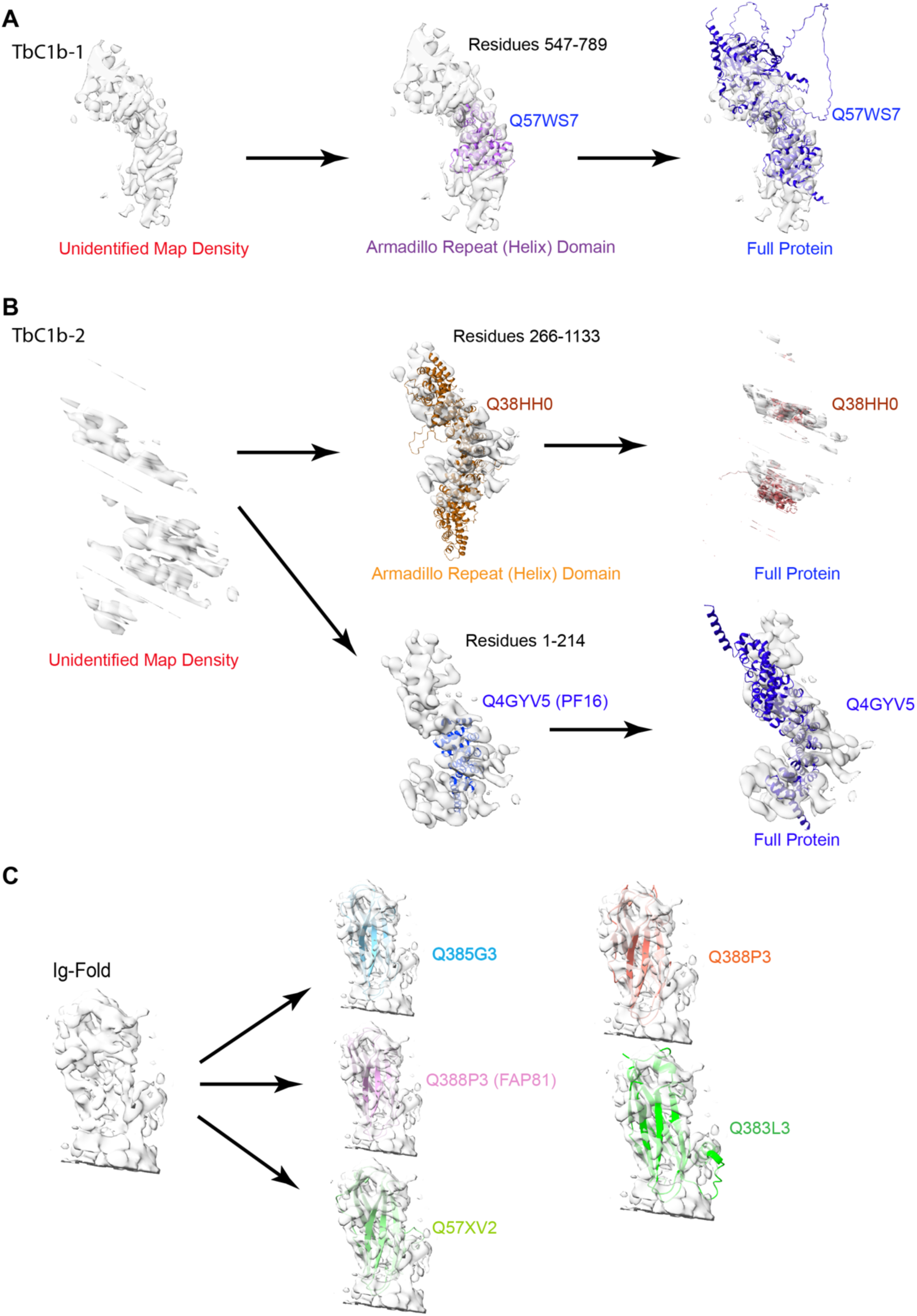
DomainFit analysis of central apparatus proteins reveals conserved folds and novel helical domains. **(A)** Domain architecture of PF16, showing armadillo repeat regions and C-terminal domains. **(B)** DomainFit-based identification of previously unannotated helical proteins and immunoglobulin-like folds within CA projections. Unmodeled densities correspond to predicted repeat domains of candidate proteins such as Q4GYV5 and Q38HH0, mapped across the C1d and C1c regions. **(C)** Protein candidate identification of Ig-fold protein by DomainFit.

## Supplementary Tables

**Supplementary Table 1:** CryoEM data collection.

|  | WT Central Pair<br>C1 Microtubule<br>(EMD-XXXXXX)<br>(PDB XXXX) | WT Central Pair<br>C2 Microtubule<br>(EMD- XXXXX)<br>(PDB XXXX) |
| --- | --- | --- |
| **Data collection and processing** |  |  |
| Magnification | 81,000 | 81,000 |
| Voltage (kV) | 300 | 300 |
| Electron exposure (e<sup>-</sup>/Å<sup>2</sup>) | 45 | 45 |
| Defocus range (µm) | -1.5 to -2.5 | -1.5 to -2.5 |
| Pixel size (Å) | 1.1 | 1.1 |
| Symmetry imposed | C1 | C1 |
| particle images (no.) | 66,294 | 49,759 |
| Map resolution (Å) | 3.8 | 4.4 |
| FSC threshold | 0.143 | 0.143 |
| Repeat unit (nm) | 16 | 16 |
| Symmetry imposed | C1 | C1 |

**Supplementary Table 2.** Central Pair Outer Proteins and MIPS: Conserved and novel Proteins in *T. brucei* central pair. Red text indicates *Chlamydomonas*-specific protein not in *T. brucei* central pair, blue text indicate *T.brucei* specific densities * indicates no atomic model built in *Chlamydomonas*)

| *T. brucei* Protein | *T. brucei* gene ID | *C. reinhardtii* | CryoEM Location | Present in proteome |
| --- | --- | --- | --- | --- |
| α-tubulin | Tb927.1.2340 | α-tubulin | Tubulin | yes |
| β-tubulin | Tb927.1.2330 | β-tubulin | Tubulin | yes |
| FAP20 | Tb927.10.2190 | FAP20 | C2a | yes |
| PF20 | Tb927.10.13960 | PF20 | Bridge C2a/e | yes |
| PF16 | Tb927.1.2670 | PF16 | Surrounding C1 | yes |
| FAP305 | Tb927.11.10350 | FAP305 (MOT17) | C1a | yes |
| FAP266/CARP10 | Tb927.11.7180 | FAP266 | C1f | yes |
| FAP65 | Tb927.10.7350 | FAP65 | C2a/e | yes |
| FAP47 | Tb927.6.620 | FAP47 | Bridge/C1 | yes |
| FAP81 | Tb927.10.13550 | FAP81 | C1e/c | yes |
| FAP101 | Tb927.10.820 | FAP101 | C1a | Only 1 of 2 replicates |
| FAP119/FAP114 | Tb927.10.5430 | FAP114/FAP119 | C1a and C1e | yes |
| FAP119 | Tb927.10.5430 | FAP114/FAP119 | C1a and C1e | yes |
| PF6 | Tb927.8.3890 | PF6 | C1a and C1e | no |
| FAP76 | Tb927.1.3250 | FAP76 | C1d/e/c | yes |
| FAP74 | Tb927.10.13550 | FAP74 | C1d/f | yes |
| Hydin | Tb927.6.3150 | Hydin | C1b/Bridge | Only 1 of 2 replicates |
| Calmodulin | Tb927.11.9790 | Calmodulin | C1a | yes |
| CPC1/ADKB | Tb927.10.830 | CPC1 | C1b | yes |
| Enolase (dimer) | Tb927.11.16410 | Enolase (Dimer) | C1b | yes |
| – | – | FAP7 | C1a | C1a |
| FAP42 | Tb927.10.11700 | FAP42 | C1b/f | yes |
| – | – | FAP46 | C1d | yes |
| – | – | FAP54 | C1d | yes |
| – | – | FAP69 | C1b | – |
| – | – | FAP194 | C1b/f | – |
| FAP99 | Tb10.v4.0044 | FAP99 | C1d | yes |
| – | – | FAP108 | C1f | – |
| – | – | FAP216 | C1e/c | – |
| FAP227 | Tb927.2.5040 | FAP227 | C1a and C1e | yes |
| FAP246/BBP68 | Tb927.10.3140 | FAP246 | C1b | yes |
| – | – | FAP275 | MIP | – |
| FAP279 | Tb927.11.5190 | FAP279 | C2a | yes |
| – | – | FAP221 | C1d/f | – |
| FAP15 (PP1c) | Tb927.11.8090 | FAP15 | C1e and C1d | yes |
| FAP70 (dimer) | Tb927.10.2380 | FAP70 | C2a | yes |
| – | – | FAP147 | C2a | no |
| – | – | FAP174 | C2a | no |
| FAP178 | Tb927.10.15280 | FAP178 | Bridge | yes |
| FAP196/Bilbo1 | Tb927.11.7560 | FAP196 | MIP | yes |
| Ig-Fold |  | – | C1a/bridge | – |
| T1bC1b-1 |  | – | C1b | – |
| T1bC1b-2 |  | – | C1b | – |
| – | – | FAP225 | MIP | – |
| – | – | FAP239 | MIP/C2a | – |
| – | – | FAP297 | C1d/f | – |
| – | – | FAP388 | MIP | – |
| – | – | FAP297 | C1d |  |
| – | – | FAP7 | C1a | – |
| – | – | FAP105 | MIP | – |
| – | – | FAP388 | MIP |  |
|  | – | FAP239 | MIP/C2a |  |
|  |  | HSP70A | C1b |  |
| – | – | FAP213 | MIP | – |
|  |  | FAP236* | – |  |
| – | – | FAP219 | Bridge | – |
|  |  | FAP424 |  |  |

